# Nanoparticle delivery of microRNA-146a regulates mechanotransduction in lung macrophages and mitigates lung injury during mechanical ventilation

**DOI:** 10.1101/796557

**Authors:** Christopher Bobba, Qinqin Fei, Vasudha Shukla, Hyunwook Lee, Pragi Patel, Rachel K Putman, Carleen Spitzer, MuChun Tsai, Mark D. Wewers, John W. Christman, Megan N. Ballinger, Samir Ghadiali, Joshua A. Englert

## Abstract

During mechanical ventilation, injurious biophysical forces exacerbate lung injury. These forces disrupt alveolar capillary barrier integrity, trigger proinflammatory mediator release, and differentially regulate genes and non-coding oligonucleotides such as microRNAs. In this study, we identify miR-146a as a mechanosensitive microRNA in alveolar macrophages that has therapeutic potential to mitigate lung injury during mechanical ventilation. We used humanized in-vitro systems, mouse models, and biospecimens from mechanically ventilated patients to elucidate the expression dynamics of miR-146a that might be required to decrease lung injury during mechanical ventilation. We found that the endogenous increase in miR-146a following injurious was relatively modest and not sufficient to prevent lung injury. However, when miR-146a was highly overexpressed using a nanoparticle-based delivery platform in vivo, it was sufficient to prevent lung injury. These data indicate that the endogenous increase in microRNA-146a during MV is a compensatory response that only partially limits VILI and that nanoparticle delivery approaches that significantly over-express microRNA-146a in AMs is an effective strategy for mitigating VILI.

## INTRODUCTION

The acute respiratory distress syndrome (ARDS) occurs in patients with pneumonia, sepsis, trauma or other pulmonary insults and is characterized by loss of alveolar-capillary barrier integrity and the development of pulmonary edema. Clinically, this manifests as impaired oxygenation, reduced lung compliance, and bilateral radiographic infiltrates on chest x-ray.(1, 2) Although supportive care with mechanical ventilation (MV) is the standard of care for ARDS patients, the physical forces generated during MV can exacerbate lung dysfunction through a phenomenon known as ventilator induced lung injury (VILI). (3, 4) VILI can also occur in mechanically ventilated patients without ARDS or those undergoing surgery.(5, 6) Injury from MV is induced through a variety of mechanisms including overdistention from high tidal volumes (volutrauma, tensile force), increased airway pressure (barotrauma, compressive force), or inadequate positive end expiratory pressure (PEEP) leading to alveolar collapse and reopening (atelectrauma, shear force). These forces directly damage the alveolar-capillary barrier, triggering the release of mediators and recruiting inflammatory cells that further damage the barrier (i.e. biotrauma).(3, 4) A more thorough understanding of the molecular mechanisms by which cells respond to mechanical forces may reveal exploitable biologic pathways that could be targeted via novel therapeutics to treat or prevent VILI.

The alveolar-capillary barrier is an extremely thin structure consisting of epithelial and endothelial cell layers. To date, most studies investigating the injurious effects of MV have focused on how these structural cells sense and respond to mechanical forces and have shown that injurious mechanical forces during volutrauma, barotrauma and atelectrauma cause plasma membrane disruption, cell detachment and activation of pro-inflammatory signaling.(7–10) In addition to structural cells, alveolar macrophages (AMs), a resident immune cell population in the lung, are known to contribute to the pathogenesis of lung injury including VILI.(11–13) AMs reside within the alveolar airspaces in close proximity to epithelial cells where they are exposed to mechanical forces during MV. Previous work has shown that macrophages are mechanosensitive and respond to cyclic stretching by increasing pro-inflammatory cytokine production.(14) However, the mechanisms by which AMs release cytokines during mechanical ventilation is not known.

MicroRNAs (miRs) are small non-coding RNA molecules that act as negative post-transcriptional regulators and play an important role in disease pathogenesis.(15) Altered miR expression plays an important role in the pathogenesis of a variety of lung diseases including: cystic fibrosis,(16) pulmonary fibrosis,(17) sarcoidosis,(18) and lung cancer.(19) Disease specific regulation makes miRs viable biomarkers, and specific miRs have been identified in acute lung injury.(20) A prior study found miRs related to innate immunity and inflammation (i.e. miR-146a and miR-155) were significantly upregulated in whole lung in response to injurious MV.(21) However, little is known about how mechanical forces during ventilation alter miR expression in specific cell types. Yehya et al (22) reported increased miR-466 expression in rat alveolar epithelial cells subjected to cyclic stretching and our lab previously demonstrated increased miR-146a expression in primary human lung epithelial cells exposed to cyclic transmural pressure.(23) We have also reported that overexpression of miR-146a reduced pressure-induced inflammation in human epithelial cells by targeting key elements of the toll-like receptor signaling pathway (IRAK1 and TRAF6)(23). However, it is not known if miR-146a is a mechanosensitive miR in AMs or if modulating miR-146a expression can be used to dampen inflammation in AMs exposed to mechanical stress during MV. In this study, we tested the hypothesis that miR-146a regulates the mechanotransduction processes in AMs that lead to VILI using both in vitro and in vivo models.

## RESULTS

### miR-146a is upregulated in primary human alveolar macrophages in response to oscillatory transmural pressure and in bronchoalveolar lavage cells from mechanically ventilated patients

To determine the mechanosensitivity of primary human AMs, AMs isolated from human donor lungs were subjected to 16 hours of oscillatory pressure at an air-liquid interface as an in vitro model of barotrauma during VILI.(23) Pressure-induced changes in pro-inflammatory cytokine secretion (Supplemental Figure 1) and miR-146a expression were measured. We focused on IL8 release in response to oscillatory pressure given that increased levels of IL8 in the circulation and bronchoalveolar lavage (BAL) are associated with worse outcomes in ARDS patients.(24, 25) Although baseline IL8 secretion varied by donor, exposure to barotrauma resulted in an increase in IL8 secretion across all donors (Figure 1A). There was also a consistent increase in miR-146a expression following oscillatory pressure in all donors (Figure 1B). We also investigated the macrophage response to oscillatory pressure using a cell line where PMA differentiated THP1 cells were exposed to oscillatory pressure as described above. These cells also exhibited increased IL8 secretion when exposed to oscillatory pressure compared to control cells not exposed to pressure (Figure 1C). Similar to primary human AMs, miR-146a expression in pressurized THP1 cells increased 2 to 4-fold (Figure 1D). To determine if force induced miR-146a is observed clinically, miR-146a was measured in BAL cells obtained from a cohort of patients that underwent a clinically indicated bronchoscopy. This cohort included spontaneously breathing patients, mechanically ventilated patients without ARDS, and mechanically ventilated patients with ARDS. Baseline characteristics of the subjects are shown in Table 1. There were no significant differences between groups in demographics. Patients undergoing mechanical ventilation had higher rates of shock and were therefore more likely to require vasopressor support. There were no differences between groups in the total number of BAL cells or differential cell counts. Although miR-146a expression was increased in mechanically ventilated patients without ARDS, levels in mechanically ventilated patients with ARDS were similar to spontaneous breathing controls (Figure 1E) raising the possibility that failure to upregulate miR-146a could exacerbate lung injury in the context of mechanical ventilation. These data indicate that miR-146a is a mechanosensitive microRNA in human AMs and BAL cells, and that AMs respond to injurious force by releasing inflammatory mediators and increasing expression of miR-146a.

**Table 1.** Patient characteristics and group analysis from Intensive Care Unit cohort*.

|  | No MV†<br>(N=15) | MV<br>(N=11) | MV + ARDS‡<br>(N=10) | Overall<br>(p-value) | No MV vs<br>MV<br>(p-value) | No MV vs<br>MV+ARDS<br>(p-value) | MV vs<br>MV+ARDS<br>(p-value) |
| --- | --- | --- | --- | --- | --- | --- | --- |
| Age (yrs)<br>mean±SD§ | 56.4±15.4 | 53.5±17.5 | 59.2±9.8 | 0.69 | 0.66 | 0.62 | 0.38 |
| Sex,<br>no. female (%) | 6 (40) | 5 (45) | 5 (50) | 0.89 | 1.0 | 0.70 | 1.0 |
| Race,<br>no. white (%) | 10 (71) | 10 (91) | 9 (90) | 0.36 | 0.34 | 0.36 | 1.0 |
| White Blood Cell<br>Count,<br>median (IQR) | 6.6<br>(2.0, 9.9) | 9.4<br>(6.9, 16.2) | 16.9<br>(11.9, 22.9) | 0.04 | 0.069 | 0.0060 | 0.11 |
| Lactate,<br>mean±SD | 1.33±0.51 | 2.14±1.23 | 2.04±1.28 | 0.42 | 0.19 | 0.28 | 0.87 |
| Vasopressors, no.<br>yes (%) | 0 | 7 (64) | 3 (30) | 0.0007 | 0.0005 | 0.052 | 0.20 |
| SIRS**, median<br>(IQR) | 1<br>(0, 3) | 2<br>(2, 3) | 4<br>(3, 4) | 0.0075 | 0.19 | 0.0054 | 0.043 |
| Systolic Blood<br>Pressure<90, no.<br>yes (%) | 0 | 9 (82) | 5 (50) | <0.0001 | <0.0001 | 0.0047 | 0.18 |
| BAL†† Cell Count<br>(median, IQR) |  |  |  |  |  |  |  |
| Total | 15 (7, 28) | 15 (3, 38) | 24 (9, 35) | 0.82 | 0.92 | 0.50 | 0.72 |
| Macrophages (%) | 29 (4, 81) | 33 (5, 70) | 43 (16, 73) | 0.98 | 0.95 | 0.90 | 0.82 |
| Neutrophils(%) | 43 (15, 79) | 62 (18, 94) | 36 (15, 72) | 0.52 | 0.56 | 0.64 | 0.23 |
| Lymphocytes (%) | 1 (0, 3) | 1 (0, 2) | 2 (0, 4) | 0.74 | 0.44 | 0.80 | 0.64 |
| Eosinophils | 0 (0, 0) | 0 (0, 0) | 0 (0, 1) | 0.16 | 0.37 | 0.058 | 0.33 |

\*Missing data – lactate: 10 in the no MV group, 1 in the MV group, and 3 in the MV+ARDS group; bronchial alveolar lavage cell counts: 2 in the no MV group, 1 in the MV group
†MV is mechanical ventilation
‡ARDS is the Acute Respiratory Distress Syndrome
§SD is standard deviation
IQR is interquartile range
\*\*SIRS is the systemic inflammatory response syndrome
††BAL is bronchial alveolar lavage

**Figure 1.**
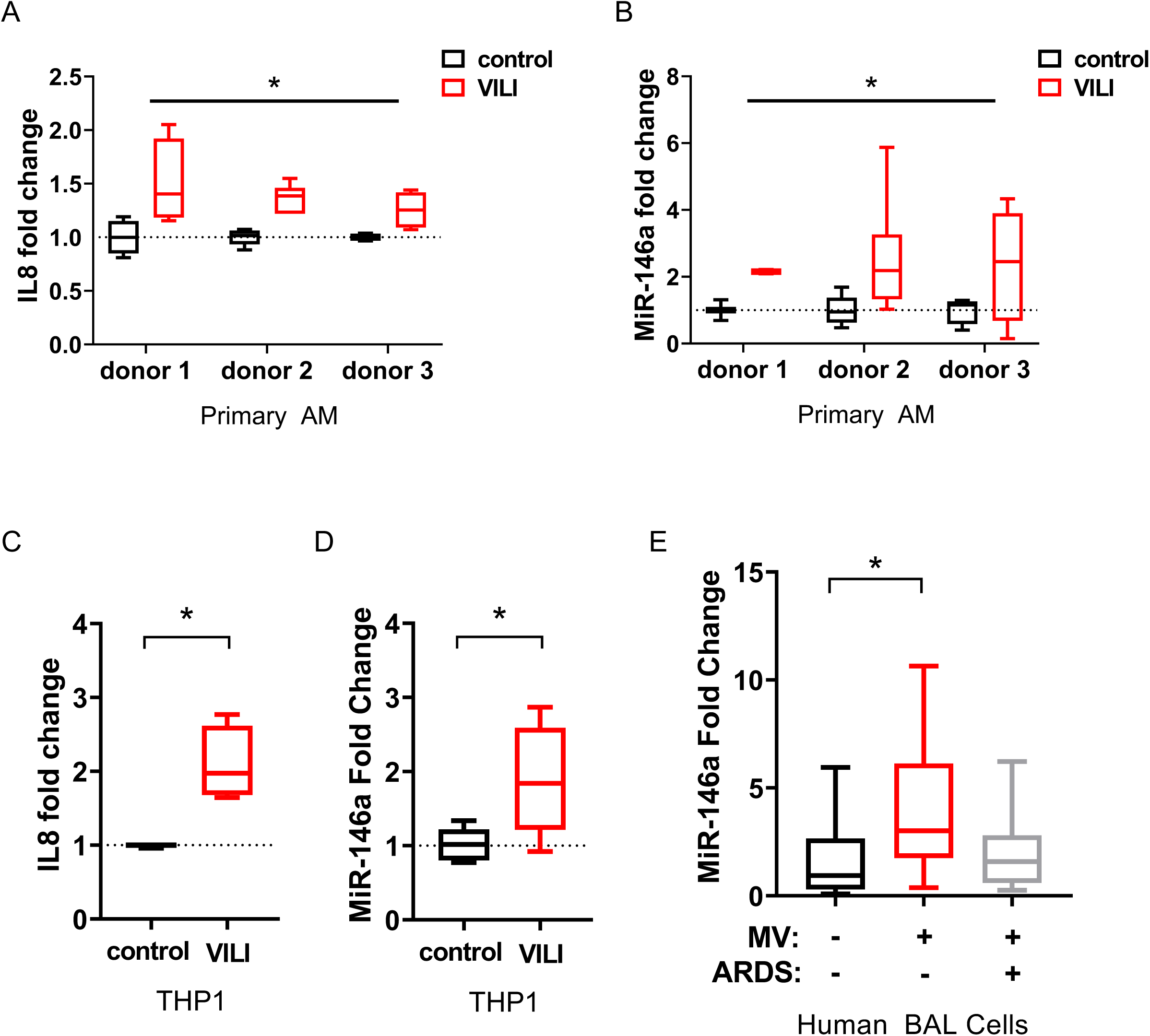
miR-146a is upregulated in primary AMs, THP1 cells, and BAL cells from an ICU cohort during VILI. **A)** Fold change in IL8 secretion in AMs from three donor lungs subjected to 16 hours of oscillatory pressure at an air-liquid interface (VILI), compared to unpressurized controls. *indicates that VILI is a statistically significant factor, p<0.0001, via 2-way ANOVA on normally distributed data, n=4 for donors 1 and 3, n=6 for donor 2. **B)** Relative miR-146a expression in AMs subjected to pressure, calculated by ΔΔCt method, normalized to unpressurized controls. *indicates that VILI is a statistically significant factor, p<0.0001, via 2-way ANOVA on normally distributed data, n=2 for donor 1, n=6 for donor 2, n=4 for donor 3. **C)** Fold change in IL8 secretion from THP1 cells subjected to 16 hours of oscillatory pressure at an air-liquid interface (VILI), compared to unpressurized controls. Data normally distributed, analyzed by student’s t-test, n=4 wells/group. **D)** Relative miR-146a expression in THP1s subjected to oscillatory pressure, calculated by ΔΔCt method, normalized to unpressurized controls. Data log-normally distributed, analyzed by student’s t-test on log_2_(fold change) n=4 wells/group. **E)** MiR-146a expression in RNA extracted from BAL cells in a cohort of patients undergoing a clinically indicated bronchoscopy. Relative expression determined by ΔΔCt method, normalized to no MV. Data log-normally distributed, analyzed by one-way ANOVA on log_2_ (fold change) with Tukey post-hoc test (n=15 for no MV group, n=11 for MV no ARDS, n=10 for MV with ARDS). * p<0.05 unless otherwise noted above

### Injurious mechanical ventilation upregulates miR-146a in AMs during murine VILI

To investigate if miR-146a expression is increased in response to injurious MV, wild-type (WT) mice were subjected to mechanical ventilation with high tidal volumes (12 cc/kg) without PEEP for 4 hours. Cells were isolated from BAL fluid and the majority of cells from mechanically ventilated and spontaneous breathing control animals were macrophages (Figure 2A, Supplemental Figure 2A-C). miR-146a expression was assessed in BAL cells and increased about 10-fold following 4 hours of VILI (Figure 2B). This is consistent with our observation of a modest increase in miR-146a expression in BAL cells from mechanically ventilated patients (Figure 1E). BAL concentration of IL6 (Figure 2C) and CXCL1/KC, the murine homolog of IL8, were increased in ventilated mice compared to spontaneous breathing controls (Figure 2D). BAL total protein, a surrogate for alveolar-capillary barrier permeability, was increased following ventilation (Figure 2E). The combination of increased inflammation and barrier dysfunction resulted in increased lung elastance (i.e. stiffness, Figure 2F) and decreased oxygenation following ventilation (Figure 2G). In summary, injurious mechanical forces during MV modestly upregulate miR-146a expression in AMs during murine VILI.

**Figure 2.**
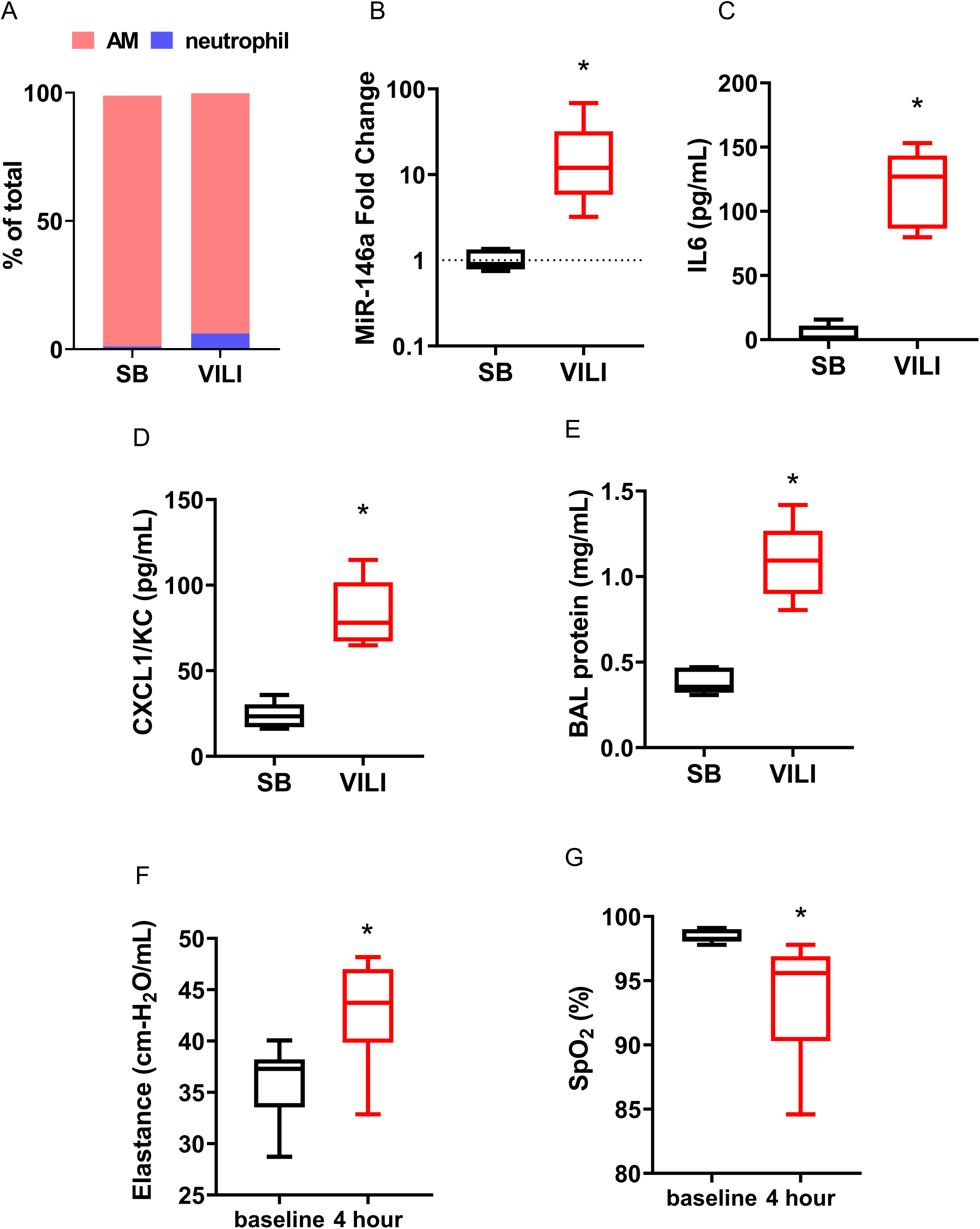
Injurious mechanical ventilation upregulates miR-146a in AMs during murine VILI. **A)** BAL differential cell counts in spontaneously breathing (SB) and ventilator induced lung injury (VILI) groups. n=5 for SB, n=6 for VILI groups. **B)** MiR-146a expression from RNA extracted from BAL cells following 4 hours injurious ventilation (VILI) or from SB controls. Relative expression determined with ΔΔCt method, normalized to SB. Data log-normally distributed, analyzed by students t-test on log_2_(fold change), n=5 for SB, n=6 for VILI groups. **C)** IL6 concentrations from SB and VILI groups. Data normally distributed, analyzed by student’s t-test. n=5 for SB, n=6 for VILI groups. **D)** KC concentration from SB and VILI groups. Data normally distributed, analyzed by student’s t-test. n=5 for SB, n=6 for VILI groups. **E)** BAL protein concentration from SB and VILI groups. Data normally distributed, analyzed by student’s t-test. n=5 for SB, n=6 for VILI groups. F) Lung tissue elastance measurements at initiation of ventilation (baseline) and at the conclusion of 4 hours ventilation. Data normally distributed, analyzed by students t-test. n=5 for SB, n=6 for VILI groups. G) Blood oxygenation saturation (SpO_2_) was measured by pulse oximetry at initiation and conclusion of ventilation. Data normally distributed, analyzed by student’s t-test. n=5 for SB, n=6 for VILI groups. * p<0.05.

### miR-146a knock-out mice have increased lung injury during mechanical ventilation

To assess whether miR-146a has an endogenous protective role in modulating VILI, miR-146a knock-out (KO) and wild-type (WT) litter-mate control mice were subjected to injurious mechanical ventilation for 4 hours and compared to spontaneous breathing (SB) control mice. There was no difference in BAL cytokine or protein concentrations in miR-146a KO mice prior to ventilation (SB in Figure 3A-C). Following ventilation, KO mice exhibited significantly increased BAL IL6 and CXCL1/KC levels (Figure 3A-B) and elevated total protein concentrations (Figure 3C). Differential cell counting revealed that global deletion of miR-146a did not significantly alter the recruitment of inflammatory cells following injurious ventilation (Figure 3D). KO mice also demonstrated a significant increase in lung elastance (Figure 3E) and decrease in oxygenation (Figure 3F) following ventilation compared to WT controls. These data demonstrate that loss of miR-146a results in increased proinflammatory cytokine secretion, disruption of the alveolar-capillary barrier, and impaired lung function compared to WT animals. Together, the data in Figures 2 and 3 indicate that miR-146a is an important negative regulator of lung injury during mechanical ventilation and suggest that the endogenous increase in miR-146a in WT mice in response to MV is not sufficient to abrogate force induced lung injury.

**Figure 3.**
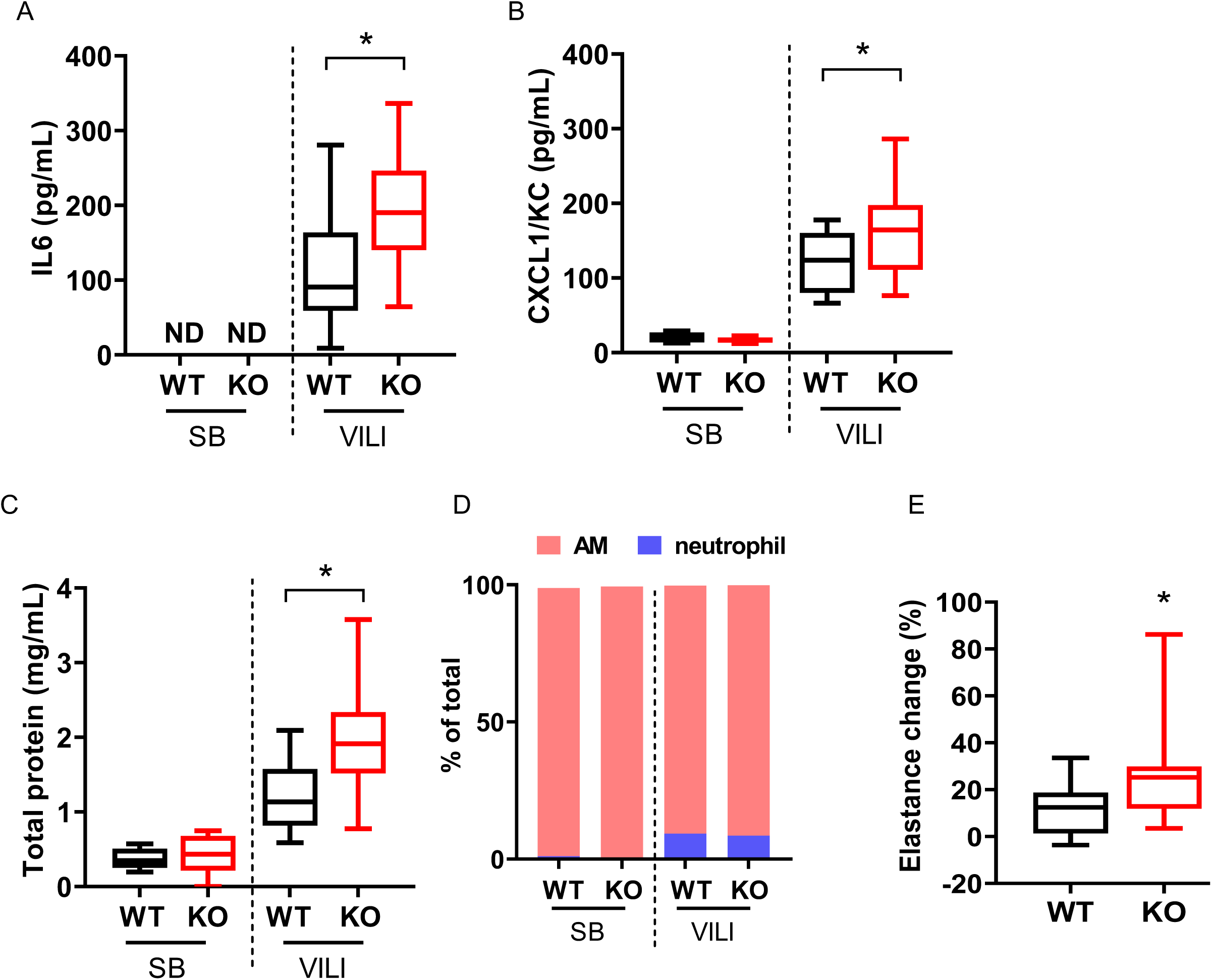
miR-146a KO mice have increased injury following mechanical ventilation. **A)** BAL IL6 from spontaneously breathing (SB) and mechanically ventilated (VILI) wild-type (WT) or miR-146a knock-out (KO) mice. Data normally distributed, analyzed by two-way ANOVA with Tukey post-hoc test. **B)** BAL KC from SB and VILI mice of either WT or KO. Data normally distributed, analyzed by two-way ANOVA with Tukey post-hoc test. **C)** BAL protein concentrations from WT and KO mice subjected to VILI (or SB controls). Data normally distributed, analyzed by two-way ANOVA with Tukey post-hoc test. **D)** BAL differential cell counts. E) Lung tissue elastance measurements during ventilation, normalized to initial values for each individual animal. Data non-normally distributed, analyzed by Mann-Whitney test. **F)** Oxygen saturation throughout duration of VILI measured via pulse oximetry. All data normally distributed, presented as mean ± SEM, analyzed by repeated measures 2-way ANOVA with Sidak’s multiple comparisons test. n=16 for WT and KO VILI groups, n=6 for WT and KO SB groups for all panels. * p<0.05.

### Endogenous expression of miR-146a in alveolar macrophages is not sufficient to mitigate VILI

A variety of cell types may be responsible for increased miR-146a expression during injurious MV in vivo. To determine the extent to which AMs contribute to VILI, mice were treated with liposomal clodronate or vehicle control prior to ventilation to deplete AMs.(26) This method led to a 50-60% reduction in the number of AMs (Supplemental Figure 3E). Consistent with a prior study in a rat model, clodronate depletion of AMs decreased lung inflammation (Supplemental Figure 3A-B), barrier permeability (Supplemental Figure 3C), and improved lung function (Supplemental Figure 3G). miR-146a levels in BAL cells were not statistically different between vehicle-control and liposomal clodronate groups, which could be due, in part, to variability between clodronate and vehicle treated animals. (Supplemental Figure 3H). However, we did observe a statistically significant correlation between miR-146a levels and the number of AMs in the BAL suggesting that AMs are an important source of miR-146a (Supplemental Figure 3I). These data also suggest that AMs play a dual role in lung injury during MV. AMs are mechanosensitive and their responses to injurious forces contribute to the development of VILI (Supplemental Figure 3A-C). In addition, these resident macrophages may also possess a compensatory pathway to mitigate lung injury by upregulating expression of miR-146a during VILI (Figures 1B,D,E and 2B).

To determine the capacity of AM-derived miR-146a to mitigate lung injury, we adoptively transferred bone marrow derived macrophages (BMDMs) from WT mice into miR-146a KO and WT control mice 6-8 hours prior to injurious MV. The adoptive transfer of WT cells into a miR-146a KO mice did not result in any change in lung injury. The miR-146a KO mice that received WT BMDMs had similar levels of BAL inflammatory cytokines and inflammatory cells (Figure 4A-B, Supplemental Figure 4, left panel), lung function (Figure 4C, left panel; Figure 4D), and barrier permeability (Figure 4E, left panel) after injurious MV as KO mice that received KO BMDMs. miR-146a levels in BAL cell pellets were modestly increased in KO mice that received WT BMDMs (Figure 4F, left panel). These data demonstrate that although there was a significant increase in miR-146a expression after adoptively transferring WT AMs into miR-146a KO mice, the level of expression was not sufficient to mitigate VILI. In a separate set of experiments, miR-146a KO AMs were adoptively transferred into WT and KO mice prior to injurious MV. Interestingly, WT mice that received miR-146a KO BMDMs had significantly higher levels of BAL IL6, KC, and neutrophils (Figures 4A-B, Supplemental Figure 4, right panel) compared to WT mice that received WT BMDMs. Despite these differences in cytokine levels and neutrophils, there were no significant differences in lung elastance (Figure 5C, right panel), oxygenation (Figure 5D), or barrier permeability (Figure 5E, right panel). Importantly, miR-146a levels were significantly decreased in the WT mice that received KO BMDMs when compared to miR-146a levels in WT mice that received WT BMDMs (Figure 4F, right panel). Data from these adoptive transfer experiments indicate that the lack of endogenous miR-146a in AMs leads to enhanced inflammation during VILI and administration of WT BMDMs is not sufficient to rescue the increased lung injury seen in miR-146a KO mice. This is likely due to the fact that the increased in miR-146a levels after adoptive transfer of WT BMDMs was modest and similar to the level observed in WT mice during ventilation (Figure 2B). We posit that this modest increase in miR-146a is insufficient to mitigate lung injury and that alternative methods to dramatically upregulate miR-146a expression may be needed to mitigate VILI.

**Figure 4.**
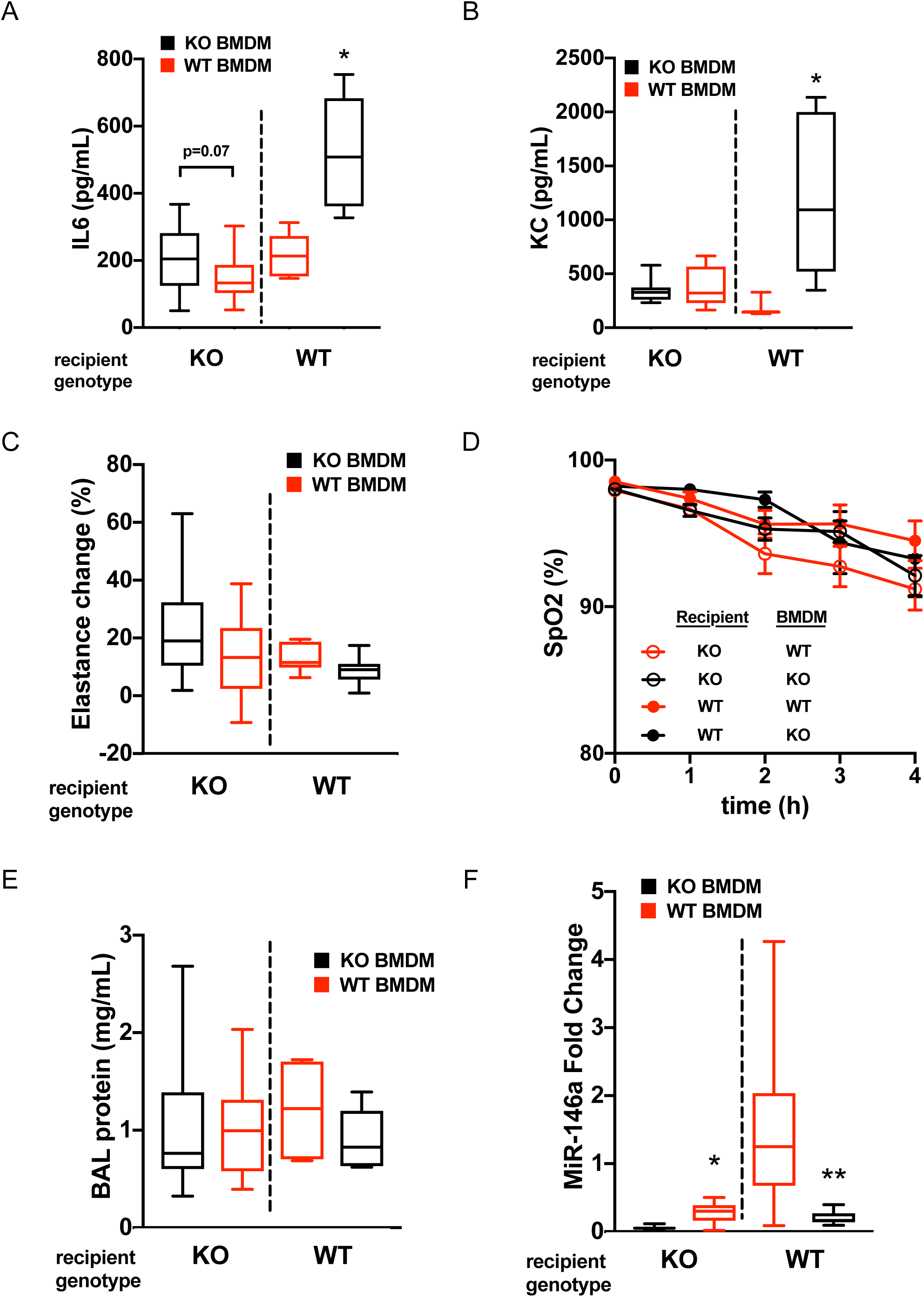
Adoptive transfer of WT macrophages into miR-146a KO mice modestly alters mIR-146a levels but does not alter VILI and adoptive transfer of KO macrophages into WT mice exacerbates lung inflammation. **A)** BAL IL6 from miR-146a KO (left panel) and WT (right panel) mice subjected to VILI following adoptive transfer of WT or miR-146a KO bone marrow derived macrophages (BMDM). All data log-normally distributed, analyzed by student’s t-test on log_2_ transformed data. For KO recipients: n=14 KO BMDM, n=13 WT BMDM. For WT recipients: n=7/group. **B)** BAL KC levels from KO (left panel) and WT (right panel) mice subjected to VILI following adoptive transfer of WT or KO BMDMs. For KO recipients: data log-normally distributed, analyzed by student’s t-test on log_2_ transformed data, n=14 KO BMDM, n=13 WT BMDM. For WT recipients: data not normally distributed, analyzed by Mann-Whitney test, n=7/group. **C)** Change in lung tissue elastance following 4 hrs MV, normalized to baseline elastance prior to MV. All data normally distributed, analyzed by student’s t-test. For KO recipients: n=14 KO BMDM, n=13 WT BMDM. For WT recipients: n=7/group. **D)** Oxygen saturation throughout duration of VILI measured via pulse oximetry. All data normally distributed, presented as mean ± SEM, analyzed by repeated measures 2 way ANOVA. For KO recipients: n=14 KO BMDM, n=13 WT BMDM. For WT recipients: n=7/group. **E)** BAL protein levels from KO (left panel) and WT (right panel) mice subjected to VILI following adoptive transfer of WT or KO BMDMs. Data log-normally distributed, analyzed by student’s t-test on log_2_ transformed data, For KO recipients: n=14 KO BMDM, n=13 WT BMDM. For WT recipients: n=7/group. **F)** miR-146a expression in BAL cell pellet RNA from KO (left panel) and WT (right panel) mice subjected to VILI following adoptive transfer of WT or KO BMDMs. Relative expression determined by ΔΔCt method, normalized to WT mice that received WT BMDMs. Data normally distributed, analyzed by student’s t-test. For KO recipients: n=14 KO BMDM, n=13 WT BMDM. For WT recipients: n=7/group. *p<0.05 compared to KO recipients that received KO BMDMs. **p<0.05 compared to WT recipients that received WT BMDMs.

**Figure 5.**
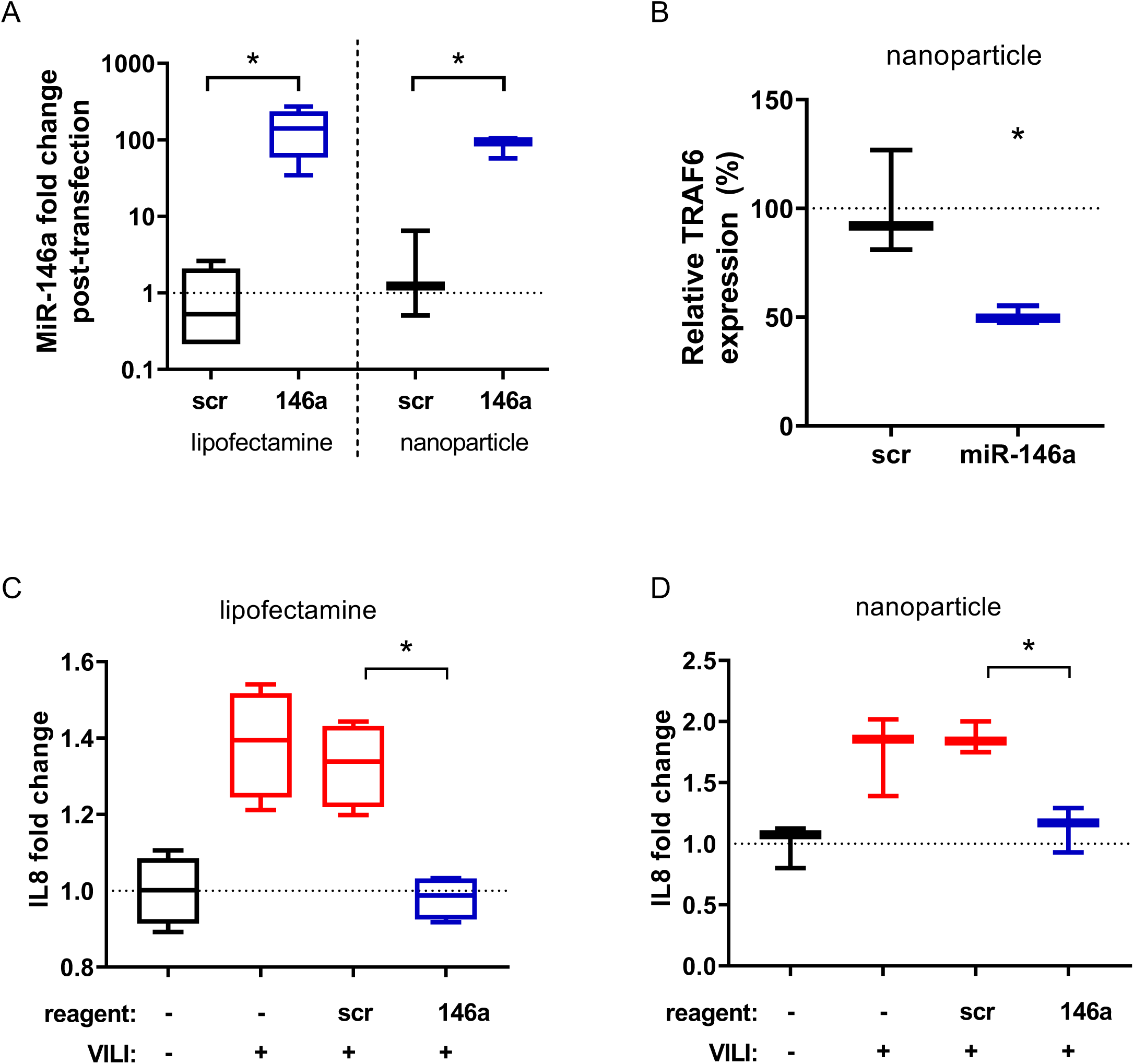
miR-146a overexpression dampens force-induced inflammation in-vitro. **A)** miR-146a expression in AMs following transient transfection with pre-miR-146a, calculated by ΔΔCt method, normalized to scramble controls. Data log-normally distributed, analyzed by student’s t-test on log_2_ fold change. n=6 per group. **B)** RelativeTRAF6 mRNA expression in AMs following nanoparticle mediated miR-146a transfection, calculated by ΔΔCt method, normalized to scramble transfected control. Data log-normally distributed, analyzed by student’s t-test on log_2_ fold change. n=3 per group. **C)** Fold change in IL8 secretion from AMs subjected to pressure alone, following scramble transfection or pre-miR-146a transfection using lipofectamine. Secretion was normalized to unpressurized control. Data normally distributed, analyzed by one-way ANOVA with post-hoc Tukey test. n=4 per group. **D)** Fold change in IL8 secretion from AMs subjected to pressure alone, following scramble transfection or pre-miR-146a transfection using custom loaded nanoparticles. Secretion normalized to unpressurized control. Data normally distributed, analyzed by one-way ANOVA with Tukey post-hoc test. n=3 per group. *p<0.05.

### miR-146a overexpression in vitro dampens the pressure-induced inflammation in primary human alveolar macrophages

To determine if miR-146a overexpression can be used to regulate force induced IL8 production, lipofectamine or nanoparticle (i.e. cationic lipoplex) based transfection techniques were used to overexpress miR-146a by 100-fold in primary human AMs (Figure 5A). miR-146a loaded nanoparticles were formulated as described in the methods and yielded particles with average an average diameter of 123nm, and near neutral zeta-potential of −0.6 mV (Supplemental Figure 5, Supplemental Table 1). Since our previous work(23) demonstrated that miR-146a regulates mechanotransduction in airway epithelial cells by targeting TRAF6, we also assessed TRAF6 expression in primary AMs following nanoparticle-based transfection. As shown in Figure 5B, TRAF6 expression decreased following miR-146a overexpression compared to scramble control. In addition, compared to cells transfected with a scrambled-miR control, primary AMs transfected with miR-146a using lipofectamine (Figure 5C) or nanoparticle-based techniques (Figure 5D) had dramatically decreased production of IL8 when subjected to our in-vitro model of barotrauma (oscillatory pressure). These data indicate that although injurious mechanical forces may result in a modest increase miR-146a expression in AMs, this is not sufficient to abrogate the pressure-induced inflammatory response. However, exogenous overexpression of miR-146a to supraphysiologic levels is capable of ameliorating pressure induced release of IL8 by primary alveolar macrophages.

### miR-146a loaded nanoparticles mitigate ventilator induced injury in vivo

To determine if overexpression of miR-146a in vivo holds therapeutic potential to reduce lung injury, we delivered pre-miR-146a loaded nanoparticles or control nanoparticles loaded with a scramble pre-miR construct to animals prior to injurious ventilation. Alveolar macrophages recovered from BAL fluid demonstrated an approximately 10,000-fold increase in miR-146a expression following delivery and ventilation, and miR-146a expression in whole lung homogenate increased about 10-to 100-fold (Figure 6A). Treatment with miR-146a nanoparticles dampened inflammation (Figure 6B-C) and reduced alveolar-capillary permeability (Figure 6D). miR-146a treated animals had improved lung compliance compared to controls (Figure 6E) and this was associated with improved blood oxygenation (Figure 6F). Consistent with our in vitro data (Figure 5B), BAL cell expression of TRAF6 was reduced following miR-146a delivery (Figure 6G). To assess whether the improvement in lung function was due to altered recruitment of inflammatory cells, we measured differential cells counts and found no difference in the number of total BAL cells (Figure 6H) or alveolar macrophages (Figure 6I). Interestingly, we did find fewer neutrophils in animals treated with miR-146a loaded nanoparticles (Figure 6J) indicating that miR-146a overexpression reduces neutrophilic lung inflammation. These findings indicate that pulmonary delivery of miR-146a loaded nanoparticles can be used to potently increase miR-146a levels and that this therapeutic overexpression mitigates lung injury during injurious mechanical ventilation.

**Figure 6.**
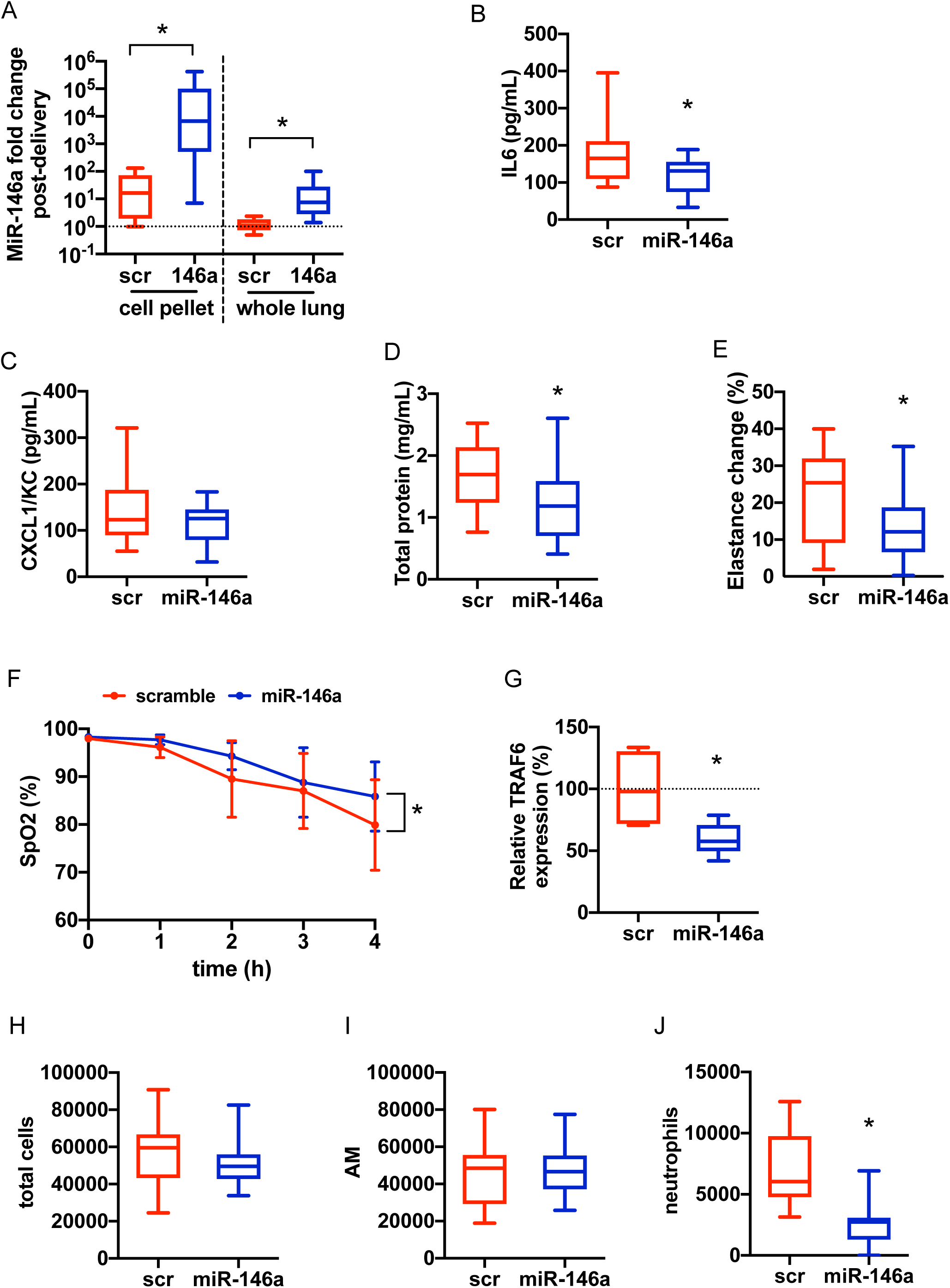
miR-146a nanoparticle delivery mitigates pro-inflammatory cytokine secretion, inflammatory cell infiltration and physiologic lung dysfunction during mechanical ventilation. **A)** MiR-146a expression in total RNA extracted from BAL cell pellet or from total RNA extracted from whole lung tissue following nanoparticle delivery and ventilation. Relative expression determined by ΔΔCt method, normalized to VILI alone. Data log-normally distributed, analyzed by student’s t-test on log_2_(fold change), between scramble and miR-146a cell pellet groups and whole lung groups independently. **B)** BAL IL6 from miR-146a loaded nanoparticle or scramble loaded nanoparticle (scr) treated mice following 4 hours injurious mechanical ventilation. Data normally distributed, analyzed by student’s t-test. **C)** BAL KC from miR-146a and scr mice following VILI. Data normally distributed, analyzed by student’s t-test. **D)** BAL protein concentration BAL from miR-146a and scr mice following VILI. All data normally distributed, analyzed by student’s t-test. **E)** Change in lung elastance during 4 hour period of ventilation. Data normally distributed, analyzed by student’s t-test. F) Oxygenation throughout duration of ventilation measured via pulse oximetry. Data normally distributed, analyzed by repeated measures ANOVA comparing miR-146a versus scramble delivery. **G)** TRAF6 transcriptional expression in total RNA extracted from BAL cell pellet, expression calculated by ΔΔCt method and normalized to scramble. Data log-normally distributed, analyzed by student’s t-test on log_2_(fold change). **H)** Total cell count obtained from BAL cell differential counts after ventilation. Data normally distributed, analyzed by student’s t-test. **I)** Alveolar macrophage (AM) cell count obtained from BAL following ventilation. Data normally distributed, analyzed by student’s t-test. **J)** Neutrophil cell count obtained from BAL differential following ventilation. Data normally distributed, analyzed by student’s t-test. n=16 per group. * p<0.05.

## DISCUSSION

Positive pressure mechanical ventilation is inherently injurious and strategies to minimize ventilator induced lung injury based on molecular pathogenesis could improve clinical outcomes. In this study we identified miR-146a as a mechanosensitive microRNA in alveolar macrophages and elucidated the expression dynamics of miR-146a that might be required to mitigate lung injury during mechanical ventilation. Primary human AMs increase pro-inflammatory cytokine secretion and miR-146a expression when exposed to injurious mechanical forces in vitro. Following MV in vivo, murine BAL cells, which are >90% AMs in our model, also increase miR-146a expression. We also demonstrated that MV in human subjects without ARDS leads to increased miR-146a expression. However, the endogenous increase in miR-146a both in vitro and in vivo is relatively small (2-10 fold) and is not sufficient to inhibit the mechanotransduction processes responsible for proinflammatory cytokine production and lung injury during ventilation. In contrast, when standard lipid-based transfection or nanoparticle-based techniques were used to significantly overexpress miR-146a in vitro by ~ 100 fold we observed significant inhibition of mechanically-induced pro-inflammatory cytokine secretion in AMs. Furthermore, our miR loaded nanoparticles effectively increased miR-146a expression in the lung in vivo (~1000-10,000 fold) and this overexpression dampened physiologic lung injury, pulmonary edema, and inflammation during injurious MV. These data indicate that miR-146a upregulation in the setting of mechanical ventilation represents an endogenous response that attempts to limit lung injury/inflammation. Although this endogenous response is insufficient to mitigate lung injury, supraphysiologic over-expression of miR-146a is a novel strategy to decrease lung injury during MV.

The concept that a modest increase in a specific microRNA is insufficient to mitigate a pro-inflammatory insult has been reported previously. For example, cigarette smoke, which induces a neutrophilic inflammatory response, was shown to increase miR-135b by ~ 20-fold while a ~ 2000-fold increase in miR-135b expression was required to suppress cigarette induced inflammation.(27) Similarly, we previously demonstrated that mechanically induced inflammation in primary human airway epithelial cells resulted in a modest 2-fold increase in miR-146a expression while a ~500 to 1000 fold increase was required to suppress mechanically-induced inflammation in these epithelial cells.(23) Therefore, the modest increase of endogenous microRNA expression in response to different pulmonary insults may represent an insufficient-compensatory response, and appropriate delivery and dosing (i.e. overexpression to appropriate levels) of microRNAs may be critical for therapeutic applications.

Although microRNAs are attractive therapeutic candidates for a wide range of diseases,(28) there are a unique set of challenges in delivering oligonucleotides to the distal lung.(29) First, although delivery to the upper airways via the oropharynx is straightforward, effective drug delivery to the distal airways and alveoli requires particle sizes on the nanometer scale.(30) Additional concerns include stability against degradation, inefficient cellular uptake, and off-target effects to other organs.(29) To overcome these challenges several solutions have been proposed, including viral vectors and lipid-based nanocarriers. Viral vectors offer excellent stability and transfection efficiency but can induce undesirable inflammatory effects. Liposomes, which are spontaneously assembled nanoparticles composed of amphiphilic lipids, that form complexes with DNA/oligonucleotides (lipoplexes) are an alternative to non-viral gene carriers. Although lipoplexes are ideal nanoparticles for drug delivery to the alveolus due to their small size, they can suffer from poor transfection efficiency if their surface is highly negatively charged.(31) Recent formulations incorporate positively charged lipids and polymers to improve uptake efficiency. We included cationic lipids in our nanoparticle formulation to neutralize their negative surface charge and capitalize on this characteristic. These lipoplexes exhibit superior distal lung delivery, excellent in vivo stability, little to no toxicity, and high transfection efficiency in vivo.(32) AMs exhibit high phagocytic capacity and are thought to be the cells in the lung that primarily uptake and process nanoparticles.(33) As a result, previous studies have used antibody-based techniques to target microRNA delivery to other cells types and limit particle uptake by AMs.(32) However, we identified Ams to be an important mechanosensitive cell type in VILI and therefore sought to deliver miR-146a to AMs directly. We therefore fabricated a non-antibody tagged multicomponent lipoplex loaded with pre-miR-146a and investigated if this delivery platform could significantly modulate miR-146a expression in the mice (Figure 6). To ensure a sufficient dose was delivered and time allowed for the pre-miR-146a construct to be processed to a mature miRNA, two doses of miR-146a encapsulated nanoparticles were given (24 hours prior and immediately prior to MV). This technique increased miR-146a expression in the BAL cells (which are primarily AMs in our murine VILI model) by 1,000 to 10,000-fold (Figure 6A) and also increased miR-146a expression in the whole lung by 10 to 100-fold. This finding indicates that lipoplex nanoparticles effectively increase miR-146a expression in AMs, and to a lesser degree increase miR-146a expression in other cell types.

Although we identified miR-146a expression in AMs as an important regulator of VILI, several other cell types also play an important role in lung injury during MV. Pulmonary microvascular endothelial cells and alveolar/airway epithelial cells also respond to elevated physical forces and may contribute to VILI.(34) Neutrophils also contribute to the inflammatory response during MV.(35) Although the role of miR-146a in small airway epithelial cells has been investigated in vitro,(23) additional studies should investigate how altering miR-146a expression in other cell types alters VILI. The current study focused on how microRNAs regulate mechanically induced inflammation during MV and how these mechanical forces can disrupt the alveolar-capillary barrier.(34) Previous studies indicate that modulating cytoskeletal structure and signaling can mitigate barrier disruption(36, 37) and future studies could investigate how microRNAs, which are known to target these pathways, modulate VILI. In vivo, AMs interact directly with the alveolar microenvironment via attachment to alveolar epithelial cells.(38) Additional studies using in vitro co-culture models could be used to investigate if this cell-cell interaction regulates mechanically-induced injury/inflammation. Lastly, although the use of primary human AMs in our study demonstrates the translational relevance of our in vitro and in vivo findings, these AMs may have different activation states, which may explain the variability in baseline cytokine expression between donors.(38) Larger scale studies using AMs isolated from a larger sample size of donor lungs could investigate the potential impact of polarization on AM contribution to VILI.

In summary, we have identified a novel pathway by which mechanotransduction in alveolar macrophage regulates VILI. In response to VILI, AMs modestly increase expression of miR-146a in an attempt to mitigate further lung injury. However, this force induced increase in miR-146a expression is insufficient to dampen lung injury and higher levels of expression are necessary to mitigate force induced injury. We have demonstrated that a novel nanoparticle-based delivery platform can be used to significantly overexpress miR-146a in vitro and in vivo and that this overexpression significantly mitigates lung injury during MV.

## METHODS

### Primary AM isolation and culture conditions

Human alveolar macrophages (AMs) or PMA differentiated THP-1 cells were used for all in vitro experiments. THP-1 cells were differentiated for 48 hours with 10nM PMA. For experiments with primary cells, AMs were isolated via ex vivo lavage from de-identified lungs rejected for transplant and obtained from Lifeline of Ohio Organ Procurements agency (Columbus, OH). All lungs were from subjects with no history of chronic lung disease or cancer and were non-smokers for at least one year. After collection, RBCs were lysed and cells were enumerated and frozen down in FBS and 10% DMSO. Prior to experiments, AMs were rapidly thawed, added to RPMI (10% FBS, 1% antibiotic/antimycotic), and centrifuged at 400g for 5 minutes. Total concentration and viability of cells was determined with trypan blue staining before seeding.

### Pressure apparatus

Oscillatory pressure was applied to AMs at an air-liquid interface using a custom design apparatus described previously.(23) Briefly, AMs were seeded on Transwell 6-well inserts and allowed to adhere for 4 hours. Media was exchanged in the basal chamber, removed from the apical chamber and the well was sealed using rubber stoppers fixed to a central channel connected to tubing. Tubes from each well were connected to a water manometer and a small animal ventilator (Harvard Apparatus, Holliston, MA, USA). Oscillatory pressure was applied for 16 hours with a 0-20 cm-H_2_O range at 0.2Hz. Media was subsequently collected for analysis by ELISA, and cellular RNA was isolated for RT-qPCR.

### Cell transfection

AMs were thawed and seeded onto Transwell inserts as described above. AMs were transfected as previously described.(39) Briefly, following adherence, AMs were transfected with Opti-MEM (with antibiotic supplement), 5nM pre-miR-146a (Thermofisher Scientific, Waltham, MA, USA), and lipofectamine (Thermofisher Scientific) solution for 24 hours. Following transfection, media was replaced with RPMI and cells were subjected to pressure as described above.

### Animal Use

All animal studies were approved by the Institutional Animal Care and Use Committee (IACUC) at Ohio State under protocols 2011A00000081-R2 and 2013A00000105-R1. MiR-146a knock-out (KO) mice(40) (stock: 016239) were purchased from the Jackson Laboratory (Bar Harbor, ME, USA) and crossed with in-house wild-type (WT) C57Bl/6J mice. KO and WT litter-mate controls were used for all experiments, with groups matched for age (age range 8-12 weeks) and sex.

### Ventilator Induced Lung Injury (VILI) Protocol

To induce VILI, mice were ventilated for 4 hours with a tidal volume (TV) of 12 cc/kg to induce volutrauma and 0 cm-H_2_O positive end expiratory pressure (PEEP) to induce atelectrauma using a Flexivent small animal ventilator (Scireq, Montreal, QC, Canada). Mice were anesthetized with ketamine and xylazine. Following induction of anesthesia, a tracheostomy cannula was placed, and mice were connected to the ventilator. Body temperature was maintained by using a heat pad beneath the animal. Baseline lung physiology measurements were obtained by performing a recruitment maneuver followed by forced oscillation to determine tissue elastance. Blood oxygenation was monitored using a mouse thigh pulse oximeter (Starr Lifesciences). At each hourly time-point the lungs were recruited (2 deep inflations) and lung physiology parameters were measured. Animals were volume resuscitated every 2 hours ventilation with 10uL/g bolus saline. At the conclusion of ventilation or spontaneous breathing (SB) controls, mice were given an overdose of anesthetic and bronchoalveolar lavage (BAL) was performed by instilling 1mL of PBS into the lungs twice and withdrawing it each time. BAL fluid was then centrifuged for 10 min at 500g and the supernatant was collected for subsequent analysis. Red blood cells were lysed with RBC lysis buffer, and differential stain was performed (Hema 3 Stat Pack, Thermofisher Scientific) according to manufacturer’s instruction. The remaining cells were resuspended in TRIzol (Qiagen, Hilden, Germany) for RNA extraction. Following BAL, lungs were harvested and snap-frozen in liquid nitrogen. For whole lung gene expression, lungs were thawed, 750uL TRIzol was added, and homogenized using a handheld tissue homogenizer (Omni International, Kennesaw GA, USA).

### Clodronate depletion

Alveolar macrophages were depleted using liposomal clodronate (Clodrosome, Brentwood, TN, USA). Mice were anesthetized with ketamine and xylazine and then administered 50uL 5mg/mL liposomal clodronate or unloaded liposomal controls via intratracheal instillation 48 hours and 24 hours prior to mechanical ventilation experiments.

### Bone-marrow-derived-macrophage (BMDM) adoptive transfer

Bone marrow derived cells were isolated from the femur and tibia of 6-8-week-old WT or miR-146a KO mice. Bone marrow derived macrophages (BMDMs) were generated by differentiating bone marrow cells for 7 days in 20ng/mL MCSF.(41) Following differentiation, BMDMs were trypsinized, counted and resuspended at 20 × 10^7^ cells/mL. 50uL of BMDM suspension (1 × 10^6^ total cells) was intratracheally instilled into mice 6-8 hours prior to ventilation.(42)

### miR-146a lipoplex fabrication, delivery, and characterization

To overexpress miR-146a in vivo, a protocol to formulate a multicomponent lipoplex was modified.(32) Briefly, empty liposomes were composed of 1,2-dioleoyl-sn-glycero-3-phosphoethanolamine (DOPE, Avanti Polar Lipids, Alabaster, AL, USA), 1,2-dioleoyl-3-trimethylammonium-propane (DOTAP, Avanti Polar Lipids), linoleic acid (LA, Sigma Aldrich, Natick, MA, USA), and D-α-Tocopherol polyethylene glycol 1000 succinate (TPGS, Sigma Aldrich). Empty liposomes were generated by mixing DOPE:DOTAP:linoleic acid:TPGS in a mole ratio of 40:10:48:2 respectively at a total concentration of 2mg/mL in HEPES buffer. Pre-miR-146a or pre-miR-scramble control (ThermoFisher Scientific) were complexed with polyethylenimine (PEI, 2000MW, Sigma Aldrich) at a 1:10 P:N (phosphate to nitrogen) ratio in HEPES buffer. Empty liposomes and miR:PEI were separately sonicated for 5 min at room temperature, then incubated for 10 min at room temperature. The two solutions were then mixed in a 1:10 nucleic acid:lipid mass ratio. MiR-loaded lipoplexes were then concentrated to 1nmol miR in 50uL by spinning at 3500g with 10K NMWL centrifugal filters (EMD Millipore, Burlington, MA, USA). 50uL of miR-loaded lipoplexes was intratracheally instilled into mice 24 hours and 0 hours before ventilation. Particle size was determined using Nanosight NS300 (Malvern, Westborough, MA, USA) and zeta potential was measured via dynamic light scattering using NanoZS (Malvern). Encapsulation efficiency was determined by (total added miRNA – unencapsulated miRNA)/(total added miRNA) x 100. Unencapsulated miRNA was determined by measuring the total miRNA in the eluent following centrifugation.

### Patient Samples

Excess BAL fluid was obtained from patients with suspected infection during a clinically indicated bronchoscopy at The Ohio State Wexner Medical Center under an approved IRB protocol (IRB 2016H0009). Potential subjects were identified by the pulmonary consult and intensive care unit teams. All patients ≥ 18 years of age were considered eligible. BAL was completed by the subject’s treating physician and samples were sent to the clinical laboratory. Once all of the ordered tests had been performed, excess BAL fluid was put on ice and transported to our laboratory. Total and differential cell counts were performed prior to centrifugation at 300g. The supernatant was aliquoted and frozen at −80C. Trizol was added to the cell pellets for future RNA isolation.

#### RNA extraction

RNA was extracted for downstream analysis by RT-qPCR using the standard phenol-chloroform RNA extraction and purification was performed with TRIzol according to manufacturer’s protocol.

#### ELISA

Enzyme linked immunosorbent assays (ELISA) were performed to assess secreted cytokine/mediator levels in media or BAL. For human cytokines, OptEIA kits (BD Biosciences, Franklin Lakes, NJ, USA) were used (IL6, IL8, IL1β, TGFβ, IL12) and the manufacturer’s protocol was followed. For mouse mediators, Duoset ELISA kits (R&D Systems, Minneapolis, MN, USA) were used (IL6, CXCL1/KC) and the manufacturer’s protocol was followed.

#### RT-qPCR

RT-qPCR with TaqMan was used to determine relative expression levels of miR-146a compared to U18 or U6 (human), sno251 (mouse) endogenous control genes (ThermoFisher, assay IDs: 478399_mir, 001204, 001093, 001236, respectively). TRAF6 relative expression was determined relative to GAPDH control (ThermoFisher, assay IDs: Hs00939742_g1 (TRAF6) Hs02786624_g1 (GAPDH) for human samples, and Mm00493836_m1 (TRAF6), Mm99999915_g1 (GADPH) for mouse). Following RNA extraction, cDNA was synthesized using high capacity cDNA transcription kit (Thermofisher). qPCR was then performed with Taqman Master Mix (Thermofisher) on Roche LightCycler480 and data was quantified by the ΔΔCT method.

#### Statistics

For laboratory-based experiments, statistical analysis was performed using Graphpad Prism 8. The distribution of all data was tested for normality via a Shapiro-Wilk test. All data are presented as box-and-whisker plots except where noted in the. Experimental sample size was determined by performing a pilot experiment with 6 animals and the sample mean and standard deviation estimates were used to calculate the final sample size with power of 0.8 and α = .05. Data were analyzed for normal or log-normal distribution. To compare two groups with normal distribution, a student’s t-test was used for comparisons or t-test on log-transformed data if data was log-normally distributed. A Mann-Whitney test was used to compare ranks if the data were not normally distributed. For analysis of qPCR data, statistical comparison was performed on log transformed fold change data. For comparison among multiple groups, all data were normally or log-normally distributed and an ANOVA was performed on log-transformed or untransformed data as appropriate. For experiments with two-independent variables, a two-way ANOVA was performed after testing data for normality as described. If a significant effect was determined by ANOVA, a Tukey post-hoc test for individual group comparisons was performed. A p-value of <0.05 was considered statistically significant. Outliers were identified using non-linear regression via the ROUT method with a Q threshold of 1%.(43) For the clinical data, association analyses between pairs of variables were conducted with Fisher’s exact tests (for categorical variables) and two-tailed t-tests or Kruskal-Wallis tests (for continuous variables as appropriate based on the normality of the data) using SAS version 9.4 (Cary, NC). Generalized linear models were used to assess for association between variables with more than two groups for comparison.

## AUTHOR CONTRIBUTIONS

CB, SNG, and JAE conceived and designed the study. CB performed the majority of experiments and analyzed the data under the supervision of SNG and JAE. Additional data were acquired and analyzed by QF, VS, HL, PP, CS, MT, MDW, and RKP. Data were interpreted by CB, MDW, JWC, MNB, SNG, and JAE. The manuscript was drafted by CB, MNB, SNG, and JAE. Critical revision of the manuscript was performed by all authors.

## ACKNOWLEDGEMENTS

This work was partially supported by NIH Grants K08 GM102695 (JAE), R56 HL142767 (JAE, SNG), R01 HL076278 (MDW) and an OSU Presidential Fellowship (CB). We would like to thank the analytical flow cytometry core for their technical assistance and the Davis Heart and Lung Research Institute at The Ohio State University.

