## Supplemental data for "Nanoparticle delivery of microRNA-146a regulates mechanotransduction in lung macrophages and mitigates lung injury during mechanical ventilation"

Running title: miR-146a reduces ventilator induced lung injury

Corresponding authors:

Joshua A. Englert, MD
Division of Pulmonary, Critical Care, & Sleep Medicine
The Ohio State University Wexner Medical Center
201 Davis Heart and Lung Research Institute
473 West 12th Avenue
Columbus, OH 43210


Samir N. Ghadiali, PhD
Department of Biomedical Engineering
The Ohio State University
402A Bevis Hall
1080 Carmack Rd.
Columbus, OH 43210


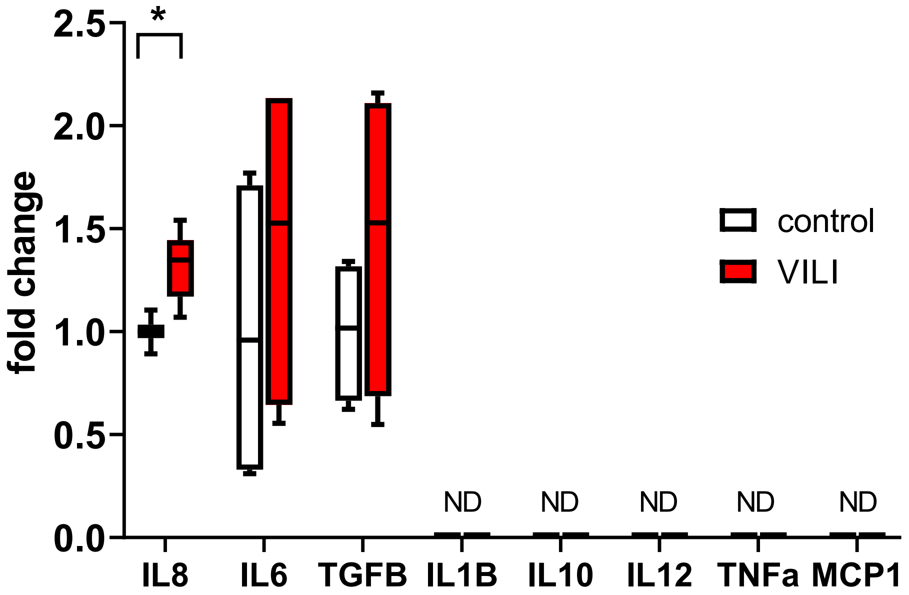


**Supplemental Figure 1. Pressure induced secretion of cytokines/mediators in primary human AMs subjected to oscillatory pressure at an air-liquid interface in vitro.** Data normally distributed, analyzed by student's t-test, n=8 per group for IL8, all others n=4. *p<.05.

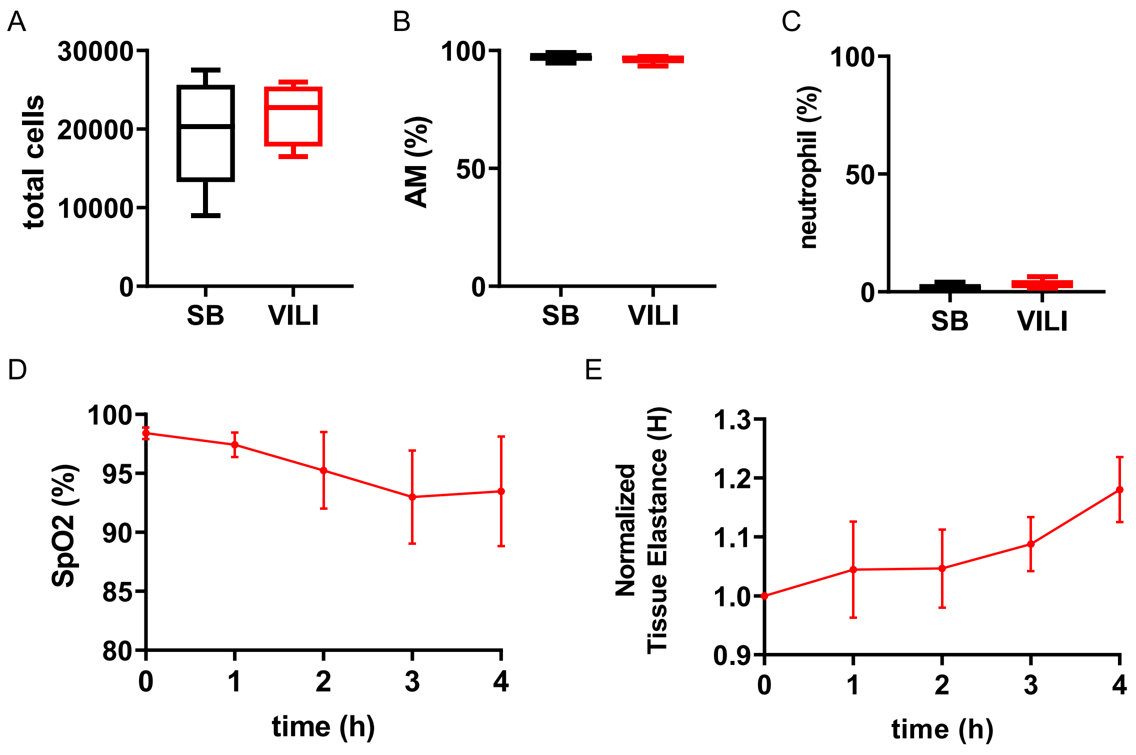


**Supplemental Figure 2. Cell differential counts in spontaneous breathing (SB) and mechanically ventilated (VILI) mice and lung physiologic data change during mechanical ventilation. A)** Total cell count obtained from BAL cell differential counts after ventilation. Data normally distributed, analyzed by student's t-test. n=5 for SB, n=6 for VILI groups. **B)** Alveolar macrophage (AM) cell count obtained from BAL following ventilation. Total cell count obtained from BAL cell differential counts after ventilation. Data normally distributed, analyzed by student's t-test. n=5 for SB, n=6 for VILI groups. **C)** Neutrophil cell count obtained from BAL differential following ventilation. Total cell count obtained from BAL cell differential counts after ventilation. Data normally distributed, analyzed by student's t-test. n=5 for SB, n=6 for VILI groups. * $p<0.05$. **D)** Tissue oxygenation throughout duration of ventilation measured via pulse oximetry. **E)** Change in lung elastance throughout 4-hour period of mechanical ventilation.

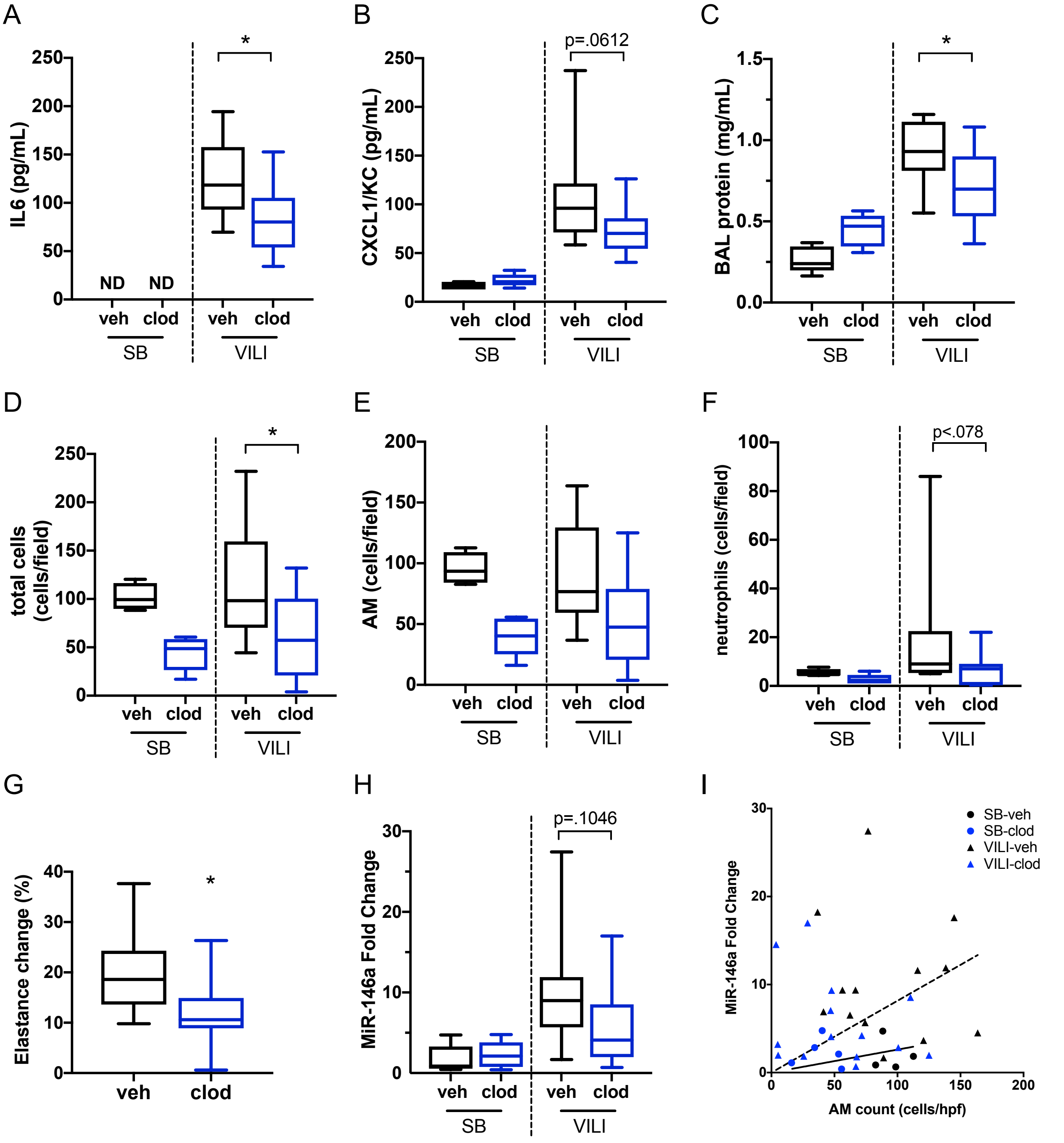


**Supplemental Figure 3. AM depletion reduces upregulation of miR-146a and dampens lung injury following injurious MV. A)** BAL IL6 from spontaneously breathing (SB) and mechanically ventilated (VILI) groups following clodronate (clod) or vehicle (veh) treatment. Data normally distributed, analyzed by two-way ANOVA with Tukey post-hoc test. n=15/group. **B)** BAL KC from SB and VILI groups following clod or veh treatments. Data log-normally distributed, analyzed by two-way ANOVA on $\log_2$ (fold change) with Tukey post-hoc test. n=14 for veh-VILI, n=15 for clod-VILI groups **C)** BAL protein concentration from SB and VILI groups following clod or veh treatment. Data normally distributed, analyzed by two-way ANOVA with Tukey post-hoc test. n=15/group. **D)** Total cell count obtained from BAL cytospin after ventilation. Data normally distributed, analyzed by two-way ANOVA with Tukey post-hoc test. n=13 for veh-VILI, n=14 for clod-VILI groups. **E)** Alveolar macrophage (AM) cell count obtained from BAL cytospin following ventilation. Data normally distributed, analyzed by two-way ANOVA with Tukey post-hoc test. n=13 for veh-VILI, n=14 for clod-VILI groups. **F)** Neutrophil cell count obtained from BAL cytospin. Data not normally distributed, analyzed by two-way ANOVA with Tukey post-hoc test. n=13 for veh-VILI, n=14 for clod-VILI groups. **G)** Change in lung tissue elastance following ventilation, normalized to baseline elastance at initiation of ventilation. Data normally distributed, analyzed by student's t-test. n=15/group. **H)** MiR-146a expression from BAL cell RNA extracted following ventilation, calculated by ΔΔCt method, normalized to veh. Data log-normally distributed, analyzed by two-way ANOVA on $\log_2$(fold change) with Tukey post-hoc test. n=15 for veh-VILI and clod-VILI groups, n=5 for veh-SB and clod-SB groups. * $p<0.05$. **I)** Correlation between AM cell count at miR-146a expression level. Solid line indicates linear regression with SB data only, $p=0.0232$, and dashed line indicates linear regression with VILI data only, $p<0.0001$.

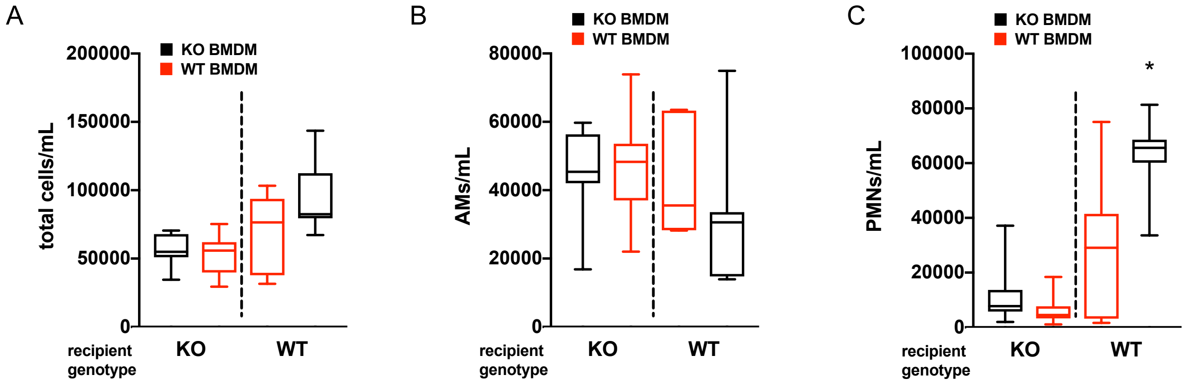


**Supplemental Figure 4. BAL total and differential cell counts from adoptive transfer experiments. A)** Total BAL cells from WT or miR-146a KO mice that received WT or KO bone marrow derived macrophages (BMDMs) prior to injurious ventilation. Counts from KO recipients were log normally distributed, analyzed by student's t-test on $\log_2$ transformed data. Counts from WT recipients were normally distributed and analyzed by student's t-test. **B)** BAL alveolar macrophages (AMs) in WT or KO recipients following injurious ventilation. Counts from KO recipients were not normally distributed, analyzed by Mann-Whitney test. Counts from WT recipients were log normally distributed and analyzed by student's t-test on $\log_2$ transformed data. **C)** BAL neutrophils (PMNs) in WT or KO recipients following injurious ventilation. KO counts were log normally distributed, analyzed by student's t-test on $\log_2$ transformed data. WT counts were normally distributed, analyzed by student's t-test. n=7/group for all WT recipients. n=14 for KO mice that received KO BMDMs and n=13 for KO mice that received WT BMDMs. *$p<0.05$

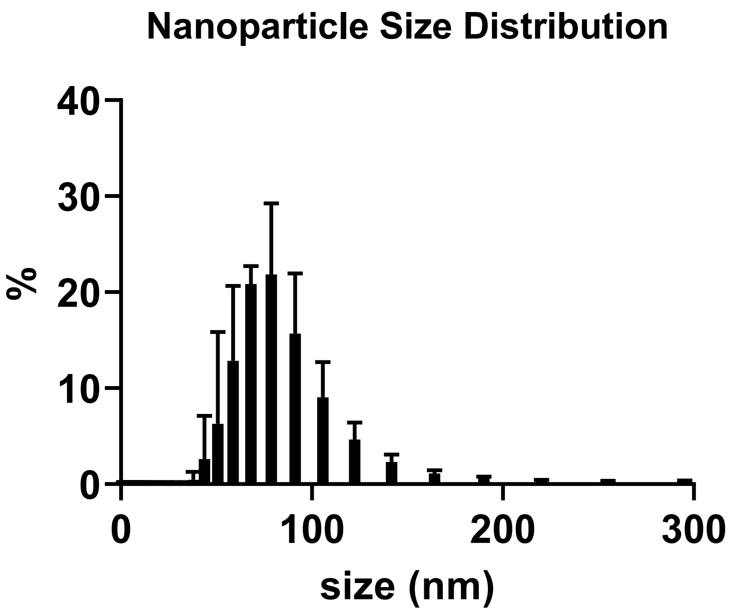


**Supplemental Figure 5. MiR-146a encapsulated nanoparticle size distribution measured by Nanosight.**

Distribution shows results averaged from 3 miR-146a encapsulated nanoparticle samples.

**Supplemental Table 1. MiR-146a encapsulated nanoparticle characteristics measured by Nanosight and dynamic light scattering.**

| | |
|---|---|
| Mean (mode)<br>Particle Diameter | 123 (105) nm |
| Mean ± SD<br>Particle Zeta Potential | -0.645 ± 1.3 mV |
| Encapsulation Efficiency | 77% |
